# Electrophysiological correlates of visual backward masking in patients with bipolar disorder

**DOI:** 10.1101/2020.05.12.090407

**Authors:** Simona Garobbio, Maya Roinishvili, Ophélie Favrod, Janir Ramos da Cruz, Eka Chkonia, Andreas Brand, Michael H. Herzog

**Affiliations:** Laboratory of Psychophysics, Brain Mind Institute, EÉcole Polytechnique Fédérale de Lausanne (EPFL), Switzerland; Laboratory of Vision Physiology, Beritashvili Centre of Experimental Biomedicine, Tbilisi, Georgia; Institute of Cognitive Neurosciences, Free University of Tbilisi, Tbilisi, Georgia; Department of Psychiatry, Tbilisi State Medical University, Tbilisi, Georgia

**Keywords:** schizophrenia, bipolar disorder, visual backward masking, endophenotype, EEG, target enhancement

## Abstract

**Background:** In visual backward masking (VBM), a target is followed by a mask that decreases target discriminability. Schizophrenia patients (SZ) show strong and reproducible masking impairments, which are associated with reduced EEG amplitudes. Patients with bipolar disorder (BP) show masking deficits, too. Here, we investigated the neural EEG correlates of VBM in BP.

**Methods:** 122 SZ, 94 unaffected controls, and 38 BP joined a standard VBM experiment. 123 SZ, 94 unaffected controls and 16 BP joined a corresponding EEG experiment, analyzed in terms of the global field power.

**Results:** As in previous studies, SZ and BP show strong masking deficits. Importantly and similarly to SZ, BP show decreased global field power amplitudes at approximately 200 ms after the target onset, compared to controls.

**Conclusions:** These results suggest that VBM deficits are not specific for schizophrenia but for a broader range of functional psychoses. Potentially, both SZ and BP show deficient target enhancement.

## 1. Introduction

Psychiatric disorders are heterogeneous and there is a considerable overlap between diseases (1). For instance, both patients with bipolar disorder (BP) and schizophrenia patients (SZ) show similar cognitive and visual deficits (2,3) as well as shared psychopathological features and genetic and psychosocial risk factors (4,5). Therefore, these two disorders, which have been traditionally considered to be distinct from each other (6,7), might belong to the same spectrum (1,8,9).

Both schizophrenia and bipolar disorder are strongly influenced by genetics. However, single-nucleotide polymorphisms (SNP) explain only a small variance of the risk for the disorders (10–12). It is therefore of great interest to find endophenotypes, which are located between the genetic and the symptomatic levels, to identify risk factors and thus improve diagnosis (13,14).

Several candidate endophenotypes have been proposed for both schizophrenia and bipolar disorder (15,16). Endophenotypes based on visual processing are of particular relevance because of their excellent reproducibility, etiology-independence, and their contributions to higher cognitive impairments such as object recognition (17–20). Visual backward masking (VBM) is such a candidate endophenotype for schizophrenia (21,22), especially the shine-through paradigm, which has a much higher sensitivity and specificity than most other perceptual and cognitive tasks (23). In backward masking, a target is followed by a mask that deteriorates performance on the target (24). In the shine-through paradigm, the target is a vertical vernier, i.e., two vertical bars slightly offset in the horizontal direction, and the mask consists of a grating of aligned verniers (see Figure 1A). Evidence for an endophenotype for schizophrenia comes from a series of studies showing that, first, SZ and schizoaffective patients have strong and reproducible performance deficits (25,26). Second, masking deficits are already present in adolescents with psychosis (27) and with first-episode psychosis (28). Third, masking deficits are state-independent (23). Fourth, healthy students scoring high in schizotypal traits also show masking deficits albeit they are highly functioning (29,30). Fifth and most importantly, unaffected siblings of SZ show masking deficits (23,31). Interestingly, siblings, adolescents with psychosis, and students scoring high in schizotypal traits are not medicated, adding further evidence that visual masking deficits are trait rather than state markers.

**Figure 1:**
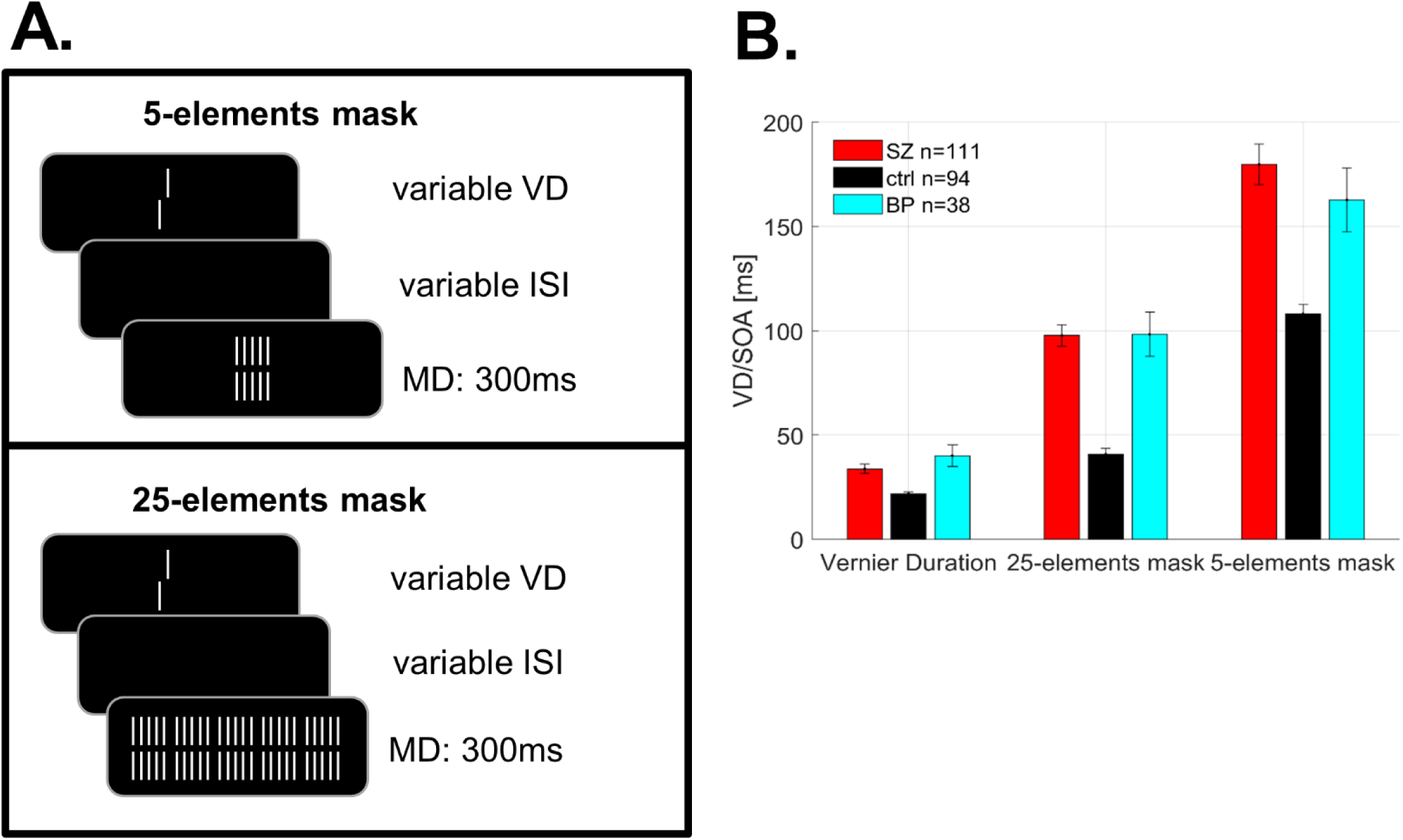
Adaptive procedure. (A) Stimulus display: The vernier duration (VD) was determined for each observer individually. Then, a mask with variable inter-stimulus interval (ISI) followed the vernier. The mask was composed of either 5- or 25-elements. The mask duration (MD) was 300 ms. (B) Behavioral results: VDs and stimulus onset asynchrony (SOA) for the two types of masks. (Note: SOA=VD+ISI, longer SOAs=stronger deficits). Mean VDs and mean SOAs of SZ (red) and BP (cyan) are higher as compared to controls (black). Error bars represent the standard error of the mean.

In SZ and in patients with first-episode psychosis, masking deficits are associated with decreased neural amplitudes of the N1 component at around 200 ms, as determined by the global field power (GFP) (28,32). Similar results were found in healthy students scoring high in schizotypal traits (30). Since BP have similar masking deficit as SZ (26), we hypothesized that they show also similar neural correlates.

## 2. Methods and Materials

### 2.1 Participants

123 SZ, 46 BP, and 94 unaffected controls participated in the experiment. Patients were recruited from the Tbilisi Mental Health Hospital. Controls were recruited from the general population in Tbilisi, aiming to match patients’ characteristics as close as possible. Participants’ age ranged from 18 to 58 years. Behavioral data of 22 out of the 46 BP were already published in a previous study (26). Also, EEG data of 110 SZ and 83 controls were published in previous work (31). Patients were diagnosed according to the Diagnostic and Statistical Manual of Mental Disorders, (DSM-IV/V) based on the Structured Clinical Interview for DSM-IV/V (Clinician Version) by an experienced psychiatrist (EC). Psychopathology of SZ was assessed by the Scales for the Assessment of Negative and Positive Symptoms (33,34). Only 31 out of the 123 SZ were assessed by the Brief Psychiatric Rating Scale (BPRS) (35). Psychopathology of all BP was assessed by the BPRS. 7 BP were diagnosed with Bipolar II disorder, and the remaining 39 with Bipolar I disorder. Most patients were medicated, only 13 SZ and 10 BP were not taking any drugs (supplemental information). General exclusion criteria were drug or alcohol dependence, severe neurological incidents or diagnoses (including head injury), developmental disorders (autism spectrum disorder or intellectual disability) or other somatic mind-altering illnesses. Family history of psychosis was an exclusion criterion for the control group. All participants had normal binocular visual acuity of at least 0.8, as measured with the Freiburg Visual Acuity Test (36). All participants were informed that they could quit the experiment at any time, and they all signed a written informed consent. All procedures complied with the Declaration of Helsinki (except for pre-registration) and were approved by the local ethics committee.

Some participants failed to perform all the tasks for various reasons (e.g., quit prematurely). 122 of the 123 SZ, all controls, and 43 out of 46 BP joined the adaptive experiment 1. We excluded 11 SZ and 5 BP because their VDs or SOAs were too long (see 2.3). All SZ and controls and 17 out of 46 BP performed the EEG experiment. 2 SZ and 1 BP were excluded due to excessive EEG artifacts (see 2.5). 122 SZ, 45 BP and 94 controls performed the WCST. 120 SZ, 45 BP and 93 controls performed the CPT. 52 SZ, 24 BP and 66 controls performed the VFT. Group characteristics and statistics are depicted in Tables 1 and 2 for the two masking experiments (adaptive and EEG), and in Table S2 in the supplemental information for the rest of the data (i.e., WCST, CPT, VFT).

**Table 1.** Group average statistics (±SD) of schizophrenia patients (SZ), patients with bipolar disorder (BP) and controls (ctrl)

|  | SZ (adaptive) | SZ (EEG) | ctrl | BP (adaptive) | BP (EEG) |
| --- | --- | --- | --- | --- | --- |
| N | 111 | 121 | 94 | 38 | 16 |
| Gender (F/M) | 17/94 | 18/103 | 47/47 | 24/14 | 11/5 |
| Age | 35.9 ± 0.8 | 36.1 ± 0.8 | 35.2 ± 0.9 | 34.4 ± 1.5 | 35.0 ± 2.2 |
| Education (years) | 13.3 ± 0.3 | 13.3 ± 0.2 | 15.2 ± 0.3 | 14.4 ± 0.4 | 14.7 ± 0.5 |
| Handedness (L/R)<sup>a</sup> | 5/106 | 6/115 | 6/88 | 0/36 | 0/13 |
| Visual acuity | 1.4 ± 0.0 | 1.4 ± 0.0 | 1.6 ± 0.0 | 1.4 ± 0.1 | 1.4 ± 0.1 |
| Illness duration (years) | 11.9 ± 0.8 | 12.1 ± 0.7 | | 10.7 ± 1.2 | 11.5 ± 2.1 |
| SANS | 10.4 ± 0.5 | 10.5 ± 0.5 | | | |
| SAPS | 9.8 ± 0.7 | 9.7 ± 0.6 | | | |
| BPRS<sup>b</sup> | 32.8 ± 0.5 | 32.6 ± 0.4 | | 31.3 ± 1.1 | 32.3 ± 1.7 |
| CPZ equivalent<sup>c</sup> | 563.7 ± 37.3 | 586.7 ± 36.8 | | 409.1 ± 62.7 | 241.3 ± 24.7 |

Abbreviations: SANS, Scale for the Assessment of Negative Symptoms; SAPS, Scale for the Assessment of Positive Symptoms; BPRS, Brief Psychiatric Rating Scale; CPZ, Chlorpromazine equivalents.
<sup>a</sup> Data from 2 BP (adaptive experiment) and 3 BP (EEG experiment) were missing.
<sup>b</sup> Only 30/111 SZ (adaptive experiment), and 30/121 SZ (EEG experiment) were considered.
<sup>c</sup> Only 98/111 SZ (adaptive experiment), 108/121 SZ (EEG experiment), 28/38 BP (adaptive experiment), and 13/16 BP (EEG experiment) were receiving medications.

**Table 2.**
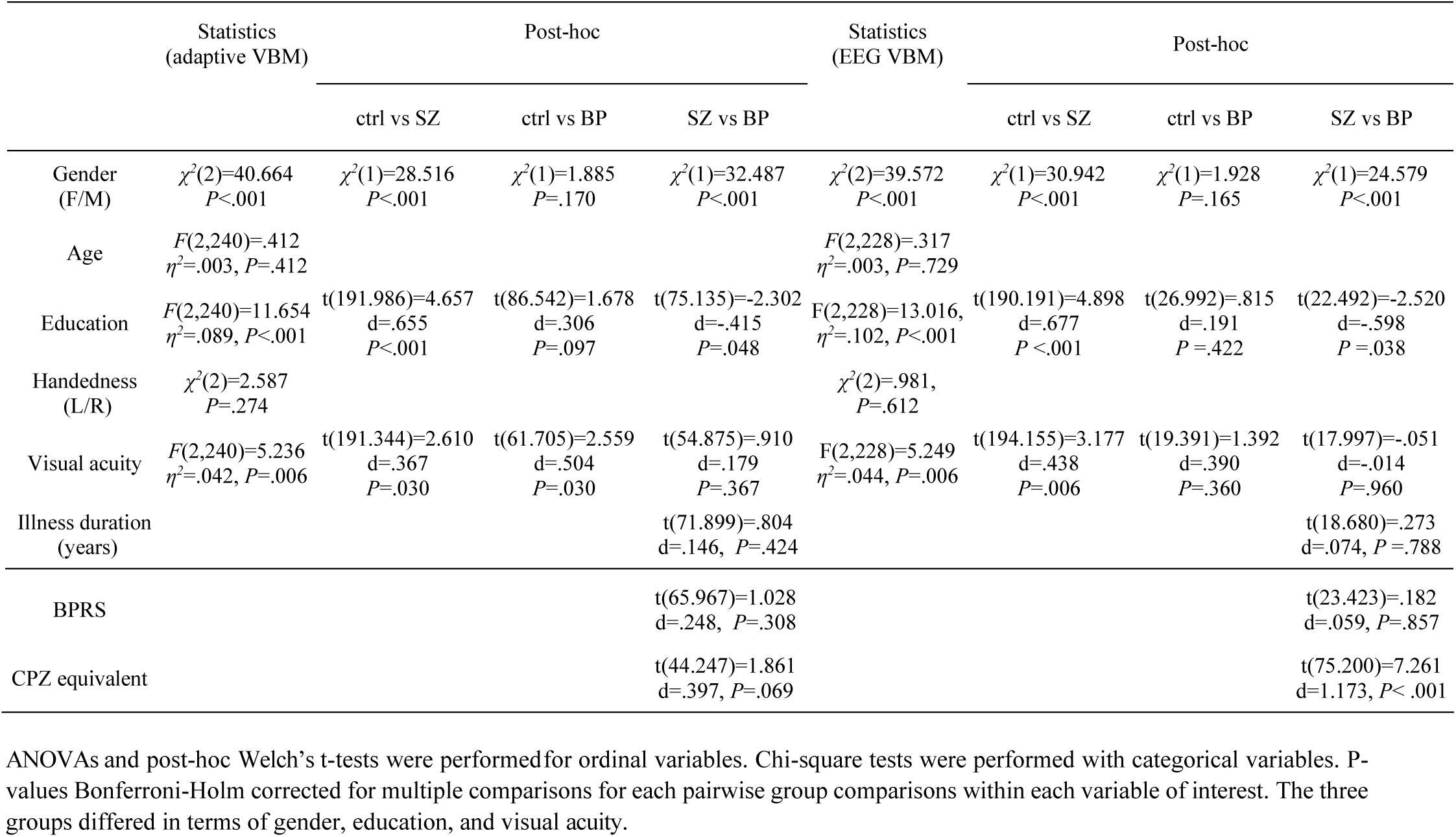
Statistical analysis of the demographical data for the visual backward masking (VBM) task (adaptive and EEG)

### 2.2 Stimuli and apparatus

Stimuli were displayed on a Siemens Fujitsu P796-1 monitor (31.0 cm (H) x 23.3 cm (V)) with a refresh rate of 100 Hz. Screen resolution was 1024×768 pixels. Participants sat in a dimly illuminated room at 3.5 m away from the monitor. At this distance, a pixel comprised about 18” (arc seconds).

The vernier stimuli were composed of two vertical bars, each 10’ (arc min) long, separated by a vertical gap of 1’. The two bars were offset in the horizontal direction of 1.2’. In each trial, the vernier offset direction was randomly chosen. The mask elements were aligned verniers, i.e., without the horizontal offset, separated horizontally by 3.3’. The vernier and the central element of the masking grating always appeared in the middle of the screen. The vernier and the two mask stimuli were presented in white (with a luminance of 100 cd/m^2^) on a black background (<1 cd/m^2^).

Participants reported the perceived offset direction of the lower bar compared to the upper bar of the vernier stimuli by hand-held button presses (left vs. right). When uncertain, participants guessed the direction. Accuracy was emphasized over speed.

### 2.3 Adaptive experiment

The detailed procedure can be found in Herzog and colleagues (25). Masking parameters were determined individually for each participant. First, an adaptive staircase procedure (PEST) (37) was used to determine the vernier duration (VD) necessary to reach 75% of correct responses for a vernier offset below 0.6’. Participants had to reach a VD shorter than 100 ms. 9 SZ and 3 BP were excluded at this stage. Second, the vernier offset was fixed to 1.2’, and individual VDs were used in the VBM task. The vernier stimulus was followed by a grating mask (lasting for 300 ms), with variable inter-stimulus interval (ISI). For each participant, the stimulus onset asynchrony (SOA = VD + ISI) to reach 75% correct responses was determined by the adaptive PEST (37). Two types of masks of either 5- or 25-elements were used (Figure 1A). Each participant performed twice; for each type of mask, the two runs were averaged and then submitted to statistical analysis. Participants with mean SOAs longer than 300 ms for the 25-elements mask and longer than 450 ms for the 5-elements mask, i.e., twice the mean SOAs of SZ in previous works (23,25,38), were excluded at this stage (2 SZ and 2 BP).

### 2.4 EEG experiment

To keep stimuli constant as required for EEG experiments, we used the same VD and SOAs for all participants. Only the 25-elements mask was used in the EEG experiment. VD was fixed to 30 ms, i.e., the average VD for SZ according to previous studies (23,25). Two SOA durations corresponding to the mean performance level of controls (30 ms) and of SZ (150 ms) were used (23,25,38).

As in previous work (31,32,38–40), the following four conditions were tested (Figure 2A): (1) Vernier Only, i.e., the vernier was presented alone for 30 ms; (2) Long SOA, i.e., the vernier was followed by the mask with an SOA of 150 ms; (3) Short SOA, i.e., the vernier was followed immediately by the mask with an SOA of 30 ms; and (4) Mask Only, i.e., the mask was presented for 300 ms (control condition). For each observer, eight blocks of 80 trials (20 trials/condition in pseudo-random order) were presented. For each recording, 160 trials per condition were computed. In the Mask Only condition, the left/right offset responses were compared to a randomly chosen notional offset.

**Figure 2:**
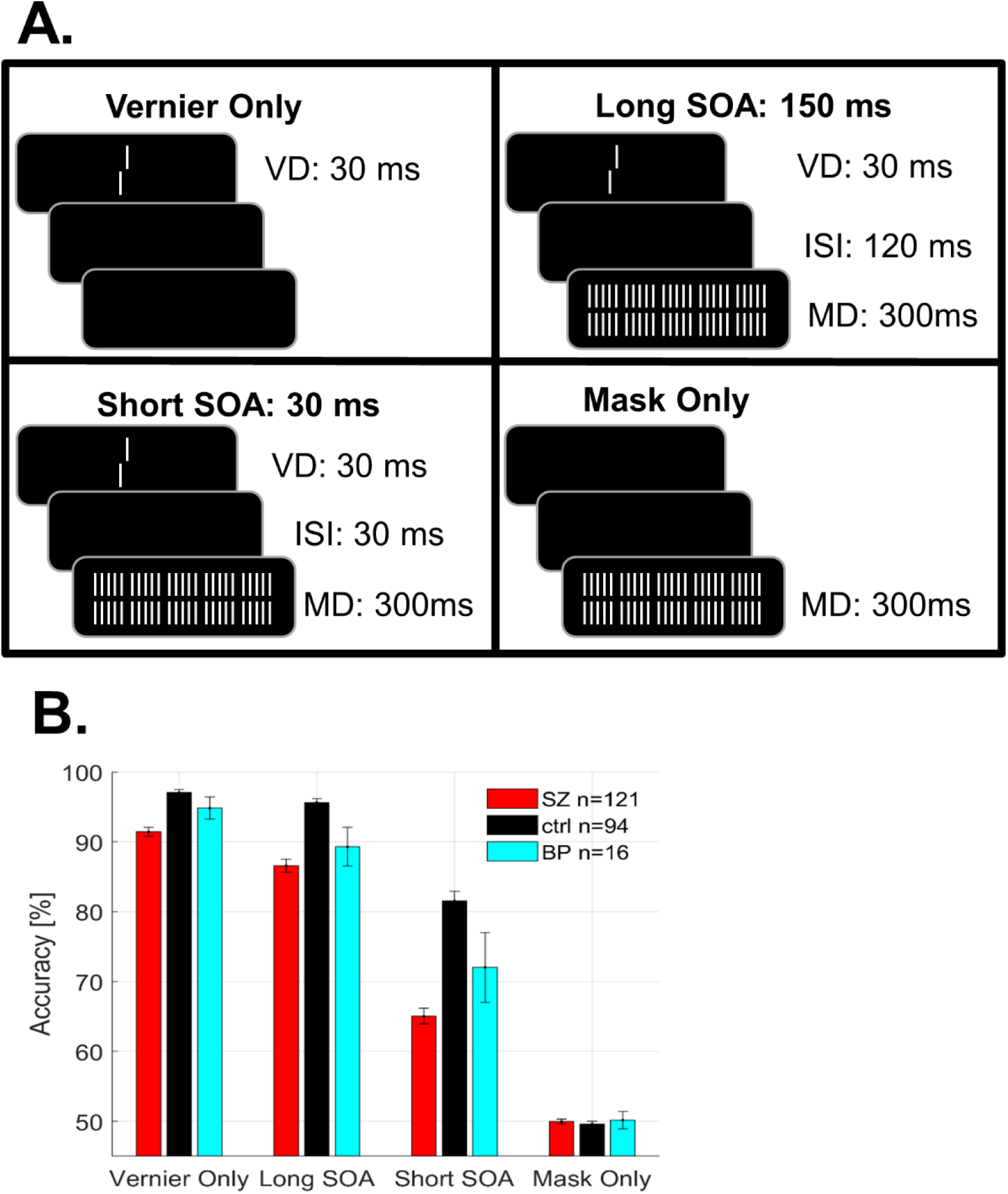
EEG experiment. (A) Stimulus display: In the Vernier Only condition, the vernier was presented alone for 30 ms. In the Short and Long SOA conditions, the vernier was followed by a mask with an SOA of either 30 or 150 ms, respectively. In the Mask Only condition, only the mask was presented. VD=vernier duration, ISI=inter-stimulus Interval, SOA=stimulus onset asynchrony, MD=mask duration, SOA=VD+ISI. (B) Accuracy for the four conditions (Vernier Only, Long SOA, Short SOA, and Mask Only). In the masking conditions, the performance of SZ (red) and BP (cyan) is lower than the one of controls (black). Error bars show the standard error of the mean.

### 2.5 EEG recording and pre-processing

EEG was recorded using a BioSemi Active Two system with 64 Ag-AgCl sintered active electrodes distributed across the scalp according to the 10/20 layout system. The sampling frequency was 2048 Hz. Offline data were pre-processed in *MATLAB* (R2012a, The MathWorks Inc., Natick, MA) using an automated pre-processing pipeline (41) (see supplemental information for details). For the EEG analysis, we excluded 1 BP due to incomplete EEG data and 2 SZ due to excessive muscular artifacts or noisy electrodes.

### 2.6 GFP analysis

The GFP was computed for each participant and each condition. The GFP is the standard deviation of potentials of all electrodes at each time point (42). The GFP is a reference-independent measure and avoids the arbitrary selection of electrodes. The GFP traces were analyzed in two ways. First, we compared GFP amplitudes of the individual evoked-related potentials (ERPs) between groups for each time point between 0 and 400 ms (205 consecutive time points), for each of the conditions separately. This analysis indicates that GFP amplitude group differences appear around the peak latencies of the GFP for each condition, and the peak latencies differ for each condition. Thus, in the second analysis we compared the GFP amplitudes at peak latencies (i.e., the N1 component) across participants and conditions. Additionally, the positive and negative components of the group grand-average ERPs were visualized by extracting the signal from two occipital electrodes (PO7 and PO8).

### 2.7 CPT, VFT, WCST

Three cognitive tests were administered: (1) The degraded continuous performance test (CPT) (43) with three blocks (720 digits, 10% targets, degradation 40%), for a total duration of 12 minutes (methodological details in Chkonia and colleagues) (44). We computed d’, which is z(hit rate)-z(false alarm rate). (2) The verbal fluency test (VFT), which was derived from the Benton controlled oral word association test (45). Participants had to report as many words as possible belonging to either the animal or fruit/vegetable category. For each category, participants had one minute to reply. The numbers of words were reported. (3) A computerized version of the Nelson test (46), which was a modified version of the Wisconsin card sorting test (WCST) (47) with 48 cards. Four measures are reported (i.e., the number of categories that subjects went through, the number of correct responses, the number of errors, and the numberof perseverative errors).

### 2.8 Statistical analysis

In the GFP timewise analysis (2.6), the GFP traces amplitudes were compared between groups for each time point trough a one-way ANOVA, and for each condition separately. The longest significant difference in the baseline (i.e., before the stimulus onset) was used as a threshold for multiple comparisons correction. Here, an effect was considered significant (*α*< .05) when at least 14 consecutive time points (about 30 ms) were significant. In doing so, short significant time intervals in the baseline or unrealistic effects (too early) were removed. This approach has been shown to partially control for multiple comparisons and false positives in EEG analyses (48–50).

Repeated measures ANOVAs were performed using JASP (https://jasp-stats.org/, version 0.11.1.0). Statistical tests were Greenhouse–Geisser corrected for violation of sphericity when necessary, and were Bonferroni-Holm corrected for multiple comparisons using RStudio (http://www.rstudio.com/, version 1.2.5033). Welch’s t-tests were used for group comparisons. Bayesian independent samples t-tests with Cauchy priors (51) were used, when opportune, to investigate non-significant group comparisons.

## 3. RESULTS

### 3.1 Behavioral results

In the adaptive experiment 1 (Figure 1B), the mean VDs and SOAs of SZ and BP are significantly longer than the one of controls, independently of the mask (5- or 25-grating mask), replicating previous results (25,26). There are no significant differences in VD and SOAs between BP and SZ. For all groups, masking is stronger with the 5-elements as compared to the 25-elements mask (Table 3).

**Table 3.**
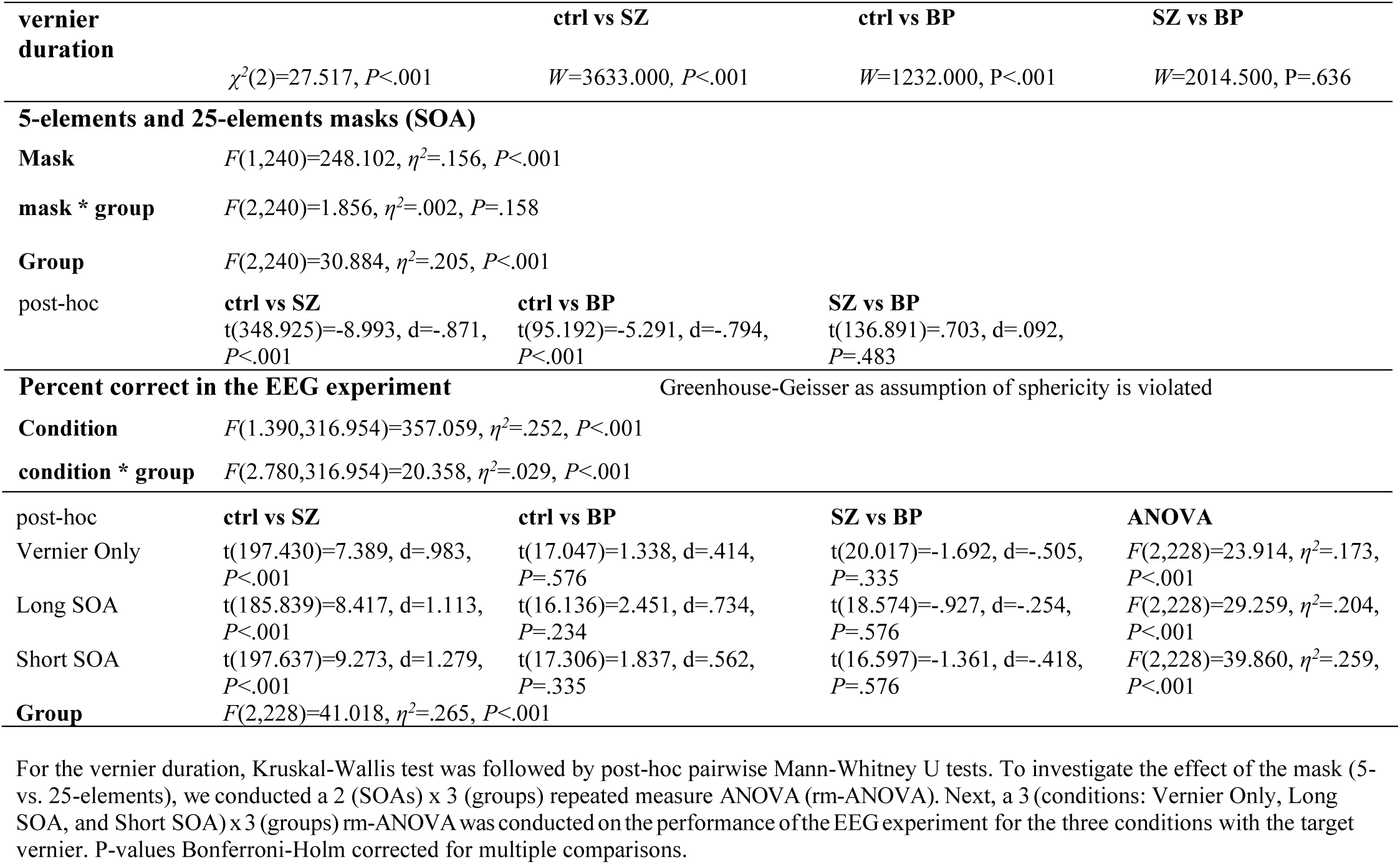
Statistical analysis for the adaptive masking experiment (VD, 5- and 25-elements masks) and the EEG experiment

In the EEG experiment (Figure 2B), we needed to use the same VD and ISI for all observers, which led to less pronounced performance differences between groups. Still, SZ performed significantly worse than controls in the three conditions with the target vernier (i.e., Vernier Only, Long SOA, Short SOA), but not for the much smaller group of the BP, most likely because of a lack of power (Table 3). A Bayesian independent samples t-tests provides weak, strong, and positive evidence (52) for the alternative hypothesis *H*_1_ (i.e., performance of controls ≠ performance of BP) for the Vernier Only (*BF*_10_=1.125), Long SOA (*BF*_10_=74.055) and Short SOA (*BF*_10_=3.242) conditions, respectively. Importantly, when only the vernier is presented the deficits are less pronounced than in the masking conditions, supporting the hypothesis that masking is the endophenotype and not the vernier duration or discrimination (53).

### 3.2 GFP analysis

The ERPs from the occipital electrodes PO7 and PO8 show a strong negative component at 200 ms after stimulus-onset, namely the N1 component (Figure 3A). Likewise for the GFP, we find a significant main effect of group around 200 ms in all conditions (Figure 3B). In the Mask Only condition, the apparent peak of BP around 280 ms is driven by one BP only. Analysis of the GFP N1 peaks amplitudes (Table 4) shows that the peaks of SZ are significantly decreased compared to controls in the three conditions with the target vernier. N1 peak amplitudes of BP are significantly lower compared to controls in the Short SOA condition only (i.e., the most challenging condition). A significant decrease is also found in the Vernier Only and Long SOA conditions, which do however not survive the correction for multiple comparisons. A Bayesian independent samples t-tests was used to compare the N1 peaks amplitudes between controls and BP for the Vernier Only and Long SOA conditions. For both conditions, according to Raftery 1995 (52), results provide weak evidence for the alternative hypothesis *H*_1_ (i.e., N1 peak controls ≠ N1 peak BP) rather than for the null hypothesis *H*_0_ (i.e., N1 peak controls = N1 peak BP), because *BF*_10_>1 (i.e., Vernier Only: *BF*_10_=2.060; Long SOA: *BF*_10_=1.797). In the Mask Only condition, GFP peak amplitudes of the three groups are comparable.

**Table 4.**
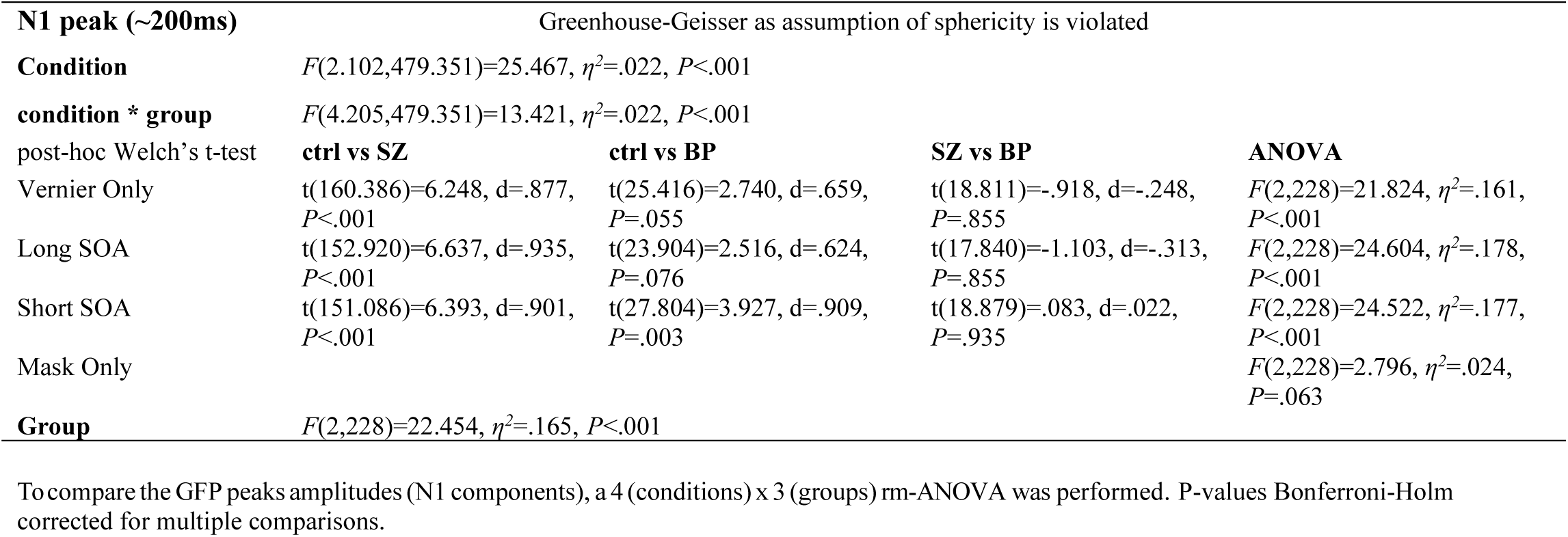
Statistical analysis for the GFP N1 peaks measured in the EEG experiment

| N1 peak (~200ms) |  | Greenhouse-Geisser as assumption of sphericity is violated |  |  |  |
| --- | --- | --- | --- | --- | --- |
| Condition | | $F(2.102, 479.351) = 25.467, \eta^2 = .022, P < .001$ | | | |
| condition * group | | $F(4.205, 479.351) = 13.421, \eta^2 = .022, P < .001$ | | | |
| post-hoc Welch's t-test |  |  |  |  |  |
|  | ctrl vs SZ | ctrl vs BP | SZ vs BP | ANOVA |  |
| Vernier Only | $t(160.386) = 6.248, d = .877, P < .001$ | $t(25.416) = 2.740, d = .659, P = .055$ | $t(18.811) = -.918, d = -.248, P = .855$ | $F(2, 228) = 21.824, \eta^2 = .161, P < .001$ | |
| Long SOA | $t(152.920) = 6.637, d = .935, P < .001$ | $t(23.904) = 2.516, d = .624, P = .076$ | $t(17.840) = -1.103, d = -.313, P = .855$ | $F(2, 228) = 24.604, \eta^2 = .178, P < .001$ | |
| Short SOA | $t(151.086) = 6.393, d = .901, P < .001$ | $t(27.804) = 3.927, d = .909, P = .003$ | $t(18.879) = .083, d = .022, P = .935$ | $F(2, 228) = 24.522, \eta^2 = .177, P < .001$ | |
| Mask Only | | | | $F(2, 228) = 2.796, \eta^2 = .024, P = .063$ | |
| Group | $F(2, 228) = 22.454, \eta^2 = .165, P < .001$ | | | | |

To compare the GFP peaks amplitudes (N1 components), a 4 (conditions) x 3 (groups) rm-ANOVA was performed. P-values Bonferroni-Holm corrected for multiple comparisons.

**Figure 3:**
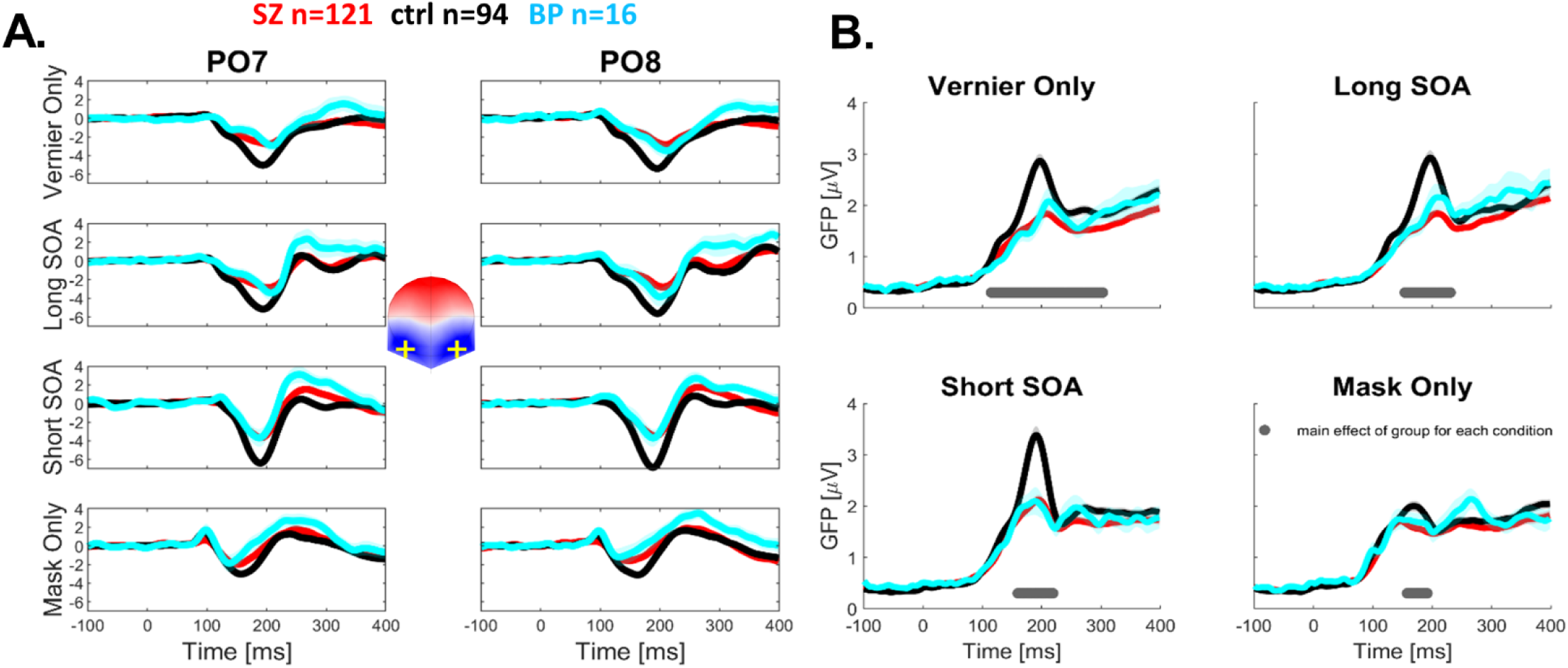
GFP analysis. (A) Group grand-average ERPs for the PO7 and PO8 electrodes. Participants showed negative deflections peaking around 200 ms, resembling a N1 component. (B) Group average global field power (GFP) time series in each condition. The bottom lines show the significant results of the timewise statistics. There is a significant difference around 200 ms in all conditions. Shaded areas indicate SEM.

### 3.3 CPT, VFT, and WCST

Overall, controls perform better than patients in all three cognitive tasks (Figure 4, Table 5). We find a significant difference between BP and controls for 5 out of 7 test variables. A lack of power may explain why we do not find a significant difference in 2 variables (i.e., VFT-animal category and WCST-errors). No significant differences were found between SZ and BP for any of the test variables. Overall, Bayesian independent samples t-tests give more evidence for H_0_ than H_1_ (i.e., SZ tests variables = BP tests variables): CPT, d: BF_01_=3.647; VFT, cat I: BF_01_=2.125, cat II: BF_01_=3.732; WCST, cat: BF_01_=3.213, corr: BF_01_=4.228, err: BF_01_=1.421, pers: BF_01_=5.360.

**Table 5.**
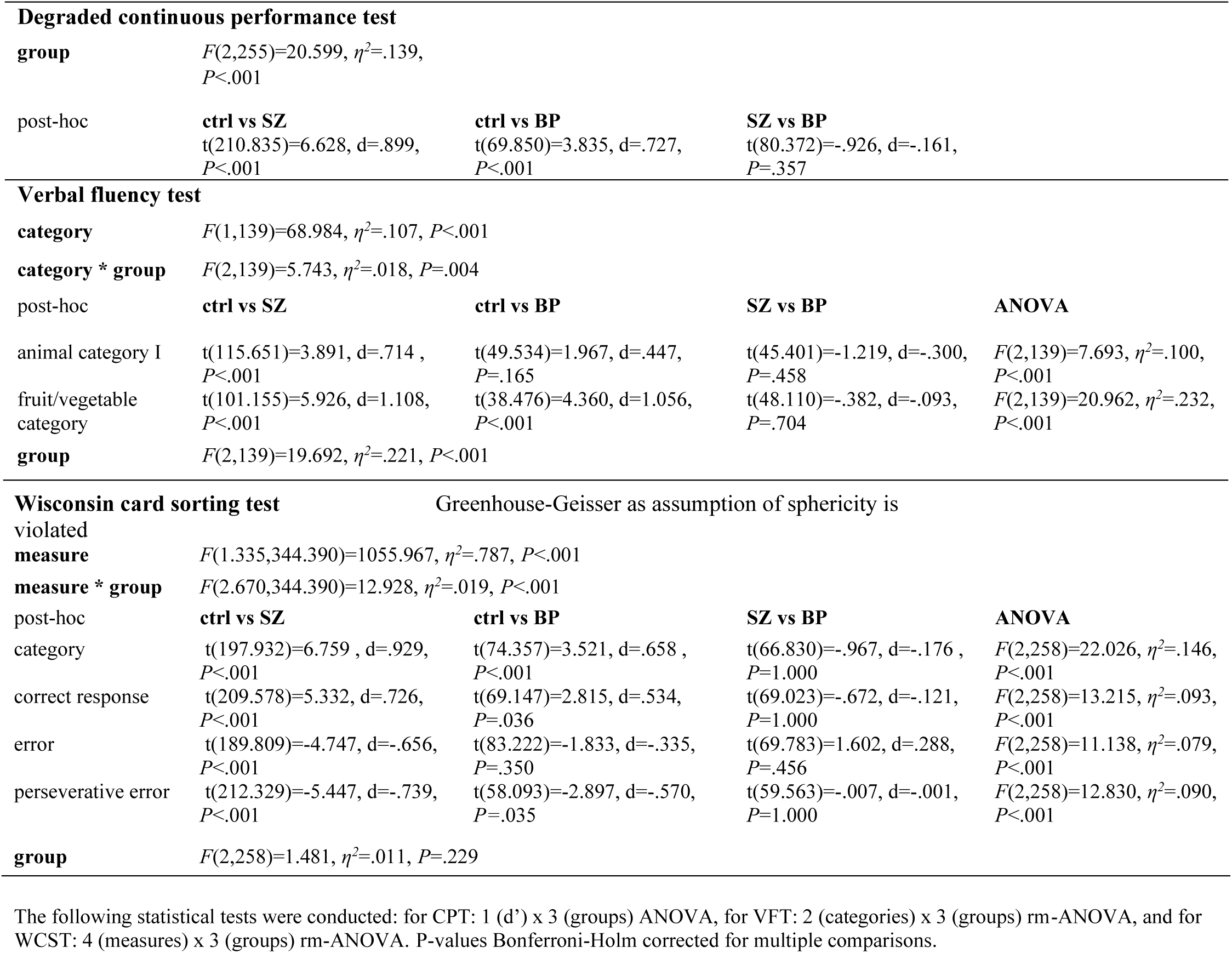
Statistical analysis for the three cognitive tasks

**Figure 4:**
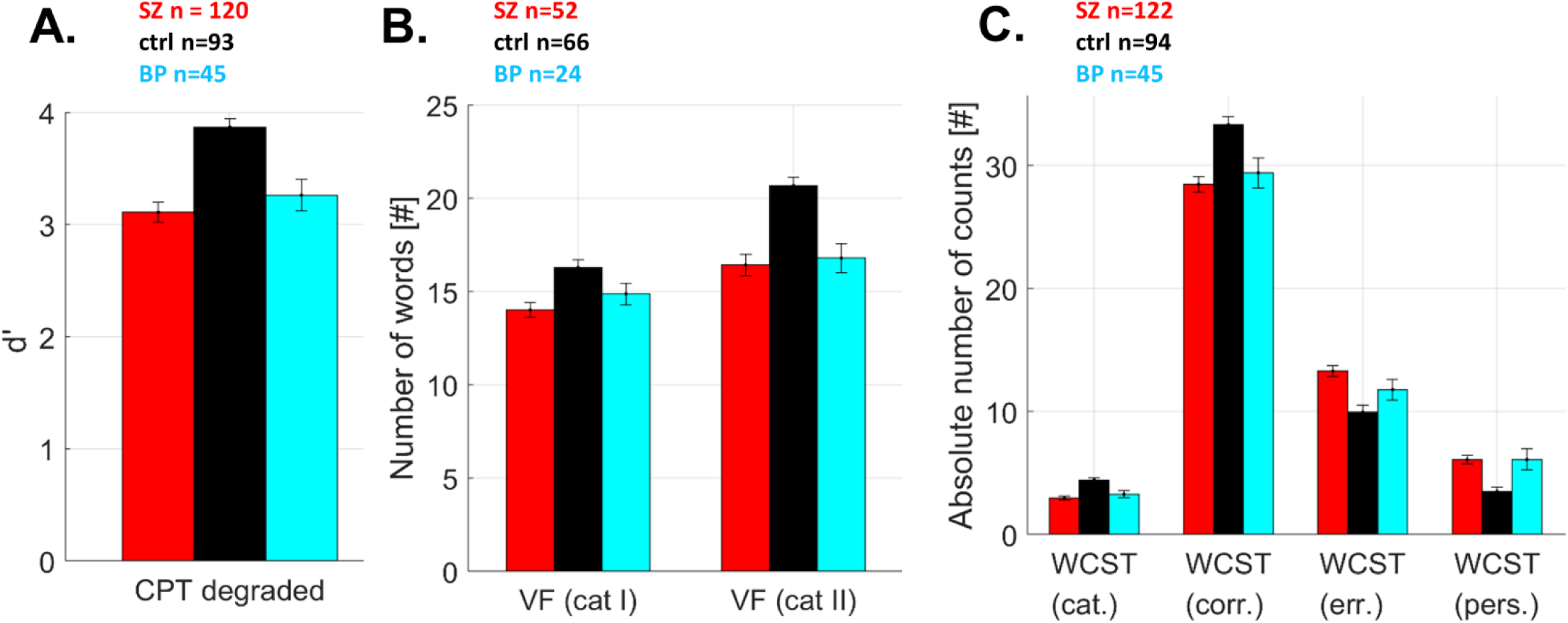
Performance for the degraded continuous performance test (CPT), the verbal fluency test (VFT), and the Wisconsin card sorting test (WCST), in the three groups (SZ, ctrl, and BP). (A) d’ for the CPT. (B) The number of words for the VFT - category I: animals, category II: vegetables/fruits. (C) The four different measures for the WCST: categories (cat.), number of correct responses (corr.), number of errors (err.), and perseverative errors (pers.). Both groups of patients performed worse than controls except for non-significant results between controls and BP for the VFT-category I and the WCST-errors.

## 4. DISCUSSION

Backward masking performance in SZ is impaired compared to unaffected controls (22,25,54–56). In BP, results are mixed. Some studies found impaired VBM performance of BP compared to controls (26,57,58), while two studies found unaffected performance of BP (59,60). However, sample sizes and hence statistical power are not large, ranging from 22 to 43 participants, which may explain the heterogeneous results. We tested 43 BP with the adaptive procedure and found that the performance of both groups of patients was strongly and similarly deteriorated compared to controls (SZ vs. controls: d=.871, BP vs. controls: d=.794). Masking deficits for BP compared to controls were also found in the EEG experiment. Our results replicate previous findings (26,57,58) and thus support the notion that schizophrenia and bipolar disorder belong to one spectrum (1).

Neurophysiologically, SZ showed strongly reduced GFP amplitudes at approximately 200 ms after the target onset compared to controls in the shine through paradigm (61). Similar results were also found in patients with first episode psychosis and students with high schizotypal traits (28,39). Here, we investigated whether the behavioral deficits found in BP are reflected neurophysiologically in a similar manner as in SZ. Qualitatively, the GFP curves of BP resemble the ones of SZ. We found significant GFP reductions of N1 peaks amplitudes in SZ and BP relative to controls in the three conditions with the target vernier. Differences between BP and controls survived the correction for multiple comparisons for the Short SOA condition (p_holm_=.003), whereas they did not for the Vernier Only (p=.011, p_holm_=.055) and the Long SOA conditions (p=.019, p_holm_=.076), likely due to a lack of statistical power. A sensitivity analysis (two-tails independent sample t-tests, alpha = 0.05, power = 80%, size BP group = 16, size control group = 94) showed that we had a sensitivity of 0.76 to detect an effect size between BP and controls, which is a large effect size according to Cohen (1988) (62). Following Bayesian analysis, we found weak evidence for a difference between BP and controls also for the Vernier Only and the Long SOA conditions. Therefore, the decreased GFP amplitudes of BP compared to controls is similar to the difference between SZ and controls and a lack of statistical power may explain why we did not find a significant difference in the Vernier Only and in the Long SOA conditions between controls and BP. No difference was found in the Mask Only condition, indicating that these deficits are specific to the target vernier and are not caused by the sheer presentation of stimuli, which may be expected by low level deficits such as generally diminished excitation.

Here, we propose the following hypothesis. The N1 amplitudes reflect, among other things, an interaction between the amplification of the target (53) and how much intrinsic effort is put in the task (40). Under normal everyday conditions, vernier-like differences go unnoticed as only a weak neural response is elicited (63). Only when the vernier is task-relevant, the human brain enhances vernier-related information to avoid overwriting by subsequently presented stimuli. Attention, recurrent processing, and neuromodulation (e.g., the cholinergic nicotinic system) may play a role in target enhancement (64–67). In SZ and BP, amplitudes are low in all target conditions. Thus, masking deficits might occur because SZ and BP cannot enhance the neural responses to the target vernier, making it more vulnerable to masking. Deficits in target enhancement happen not only in vision but also in other sensory modalities, as reflected in the mismatch negativity, auditory P3, and P50 suppression in both SZ and BP (68–70). Siblings of SZ exhibited masking behavioral deficits but surprisingly higher GFP N1 peak amplitudes compared to controls, suggesting a compensation mechanism (31). Siblings of SZ may engage more effort allowing them to recruit more neural resources to partially compensate for their behavioral deficits, if the task is not too challenging. Very preliminary results (n=4) with siblings of BP go in the same direction (supplementary Figure S1). Depressive patients showed no behavioral deficits but their N1 peaks amplitudes were reduced, though not at the level of SZ (40). This suggests that depressive patients can stabilize the neural representation of the target, making it less prone to masking and that their low amplitudes might represent less intrinsic effort.

In DSM (7) bipolar disorder and depression are thought to belong to the same family of affective disorders. However, our results show that in terms of neurocognitive performance (VBM) and the underlying brain processes (EEG), bipolar disorder is more similar to schizophrenia than to depression.

Regarding the CPT, VFT, and WCST, controls performed better than patients in all tasks in agreement with previous studies (71,72). We found a significant difference between BP and controls for 5 out of 7 output measures (Table 5). A lack of power may explain why we did not find a significant difference in all measures.

Limitations. First, sample size of the BP group in the EEG experiment was small compared to the two other groups. Thus, the lack of a clear statistical difference of the N1 amplitude between BP and controls might be due to the small sample size of BP. Second, the three groups differed in terms of gender, education, and visual acuity. To control for these variables, which have inconsistently shown to play a role in VBM performance (73), a supplementary statistical analysis including gender as a factor and visual acuity and education as covariates was conducted. The analysis showed that, overall, results were comparable to the ones obtained in the main analysis (results are shown and discussed in supplementary Tables S4, S5, and S6). Third, severity of the disorder and medications can introduce confounding factors (74–76). Generally, severity (BPRS) did not correlate with masking outcomes, in particular in BP (supplementary Table S7, left). Contrary to SZ (31), in the bipolar group, there was no correlation of CPZ and performance or N1 amplitudes (supplementary Table S7, right). However, we did not consider mood stabilizing medication. One way to bypass these confounds is to test unaffected siblings of BP. Bipolar disorder has a high heritability (70%-85%) (77) and brothers and sisters of BP, similar to siblings of SZ, have an empirical risk of approximately 10-fold higher to develop the disorder than the general population (78,79). So far, no deficits were found in siblings of BP in the literature (80,81). Finally, similar deficits in masking performance and EEG do not guarantee similar causes.

In summary, we found that BP show similar masking and EEG abnormalities as SZ, suggesting that similar mechanisms are at work.

## Acknowledgments

This work was funded by the National Centre of Competence in Research (NCCR) Synapsy financed by the Swiss National Science Foundation under grant 51NF40-185897 and by the “Knowledge Foundation” of Georgia.

BioRxiv preprint posted online May 12, 2020.

