## Supplemental Information for "Electrophysiological correlates of visual backward masking in patients with bipolar disorder"

**Affiliations:**

**Running title**: EEG correlates of VBM in patients with bipolar disorder

**Keywords:** schizophrenia; bipolar disorder; visual backward masking; endophenotype; EEG; target enhancement

### Participants: medication taken by patients with both bipolar disorder and schizophrenia

Schizophrenia patients received as antipsychotic medication clozapine, haloperidol, risperidone, perazine, trifluoperazine, fluphenazine, levomepromazine, chlorprothixene, zuclopenthixol, amisulpride, quetiapine, most patients took more than one drug. 74 patients took trihexyphenidyl against parkinsonian side effects, 7 received mood stabilizers (valproate, carbamazepin, lamotrigine), 12 patients took amitryptiline, and 12 took benzodiazepines. Bipolar patients (BP) took neuroleptics (olanzapine, levomepromazine, chlorprothixene, haloperidol, clozapine, zuclopenthixol, quetiapine, amisulpride, thioridazine, trifluoperazine, risperidone, aripiprazole), mood stabilizers (valproate, carbamazepine, lamotrigine, lithium) and antidepressants (fluvoxamine, trazodone, amitryptiline, venlafaxine, sertraline, buspirone). 10 patients took trihexyphenidy, l against parkinsonian side effects, 5 BP took benzodiazepines.

### EEG recording and pre-processing: automatic pre-processing pipeline

The automatic pre-processing pipeline (APP) (1) included the following steps: filtering via a bandpass filter of 1-40 Hz; removal of line-noise; re-referencing to the biweight estimate of the average of all electrodes; removal and 3D spline interpolation of bad electrodes; removal of bad epochs; independent component analysis (ICA) to remove artifacts related to eye movements, muscle activity, and bad electrodes; and removal of epoch artifacts. The clean EEG data were then re-referenced to the common average reference. We extracted EEG epochs from 100 ms before (baseline) to 400 ms after stimulus onset. The averaged epochs for each participant were baseline corrected. On average, 0.5 channels per EEG recording were interpolated. The proportion of rejected trials was about 2% for each EEG recording.

**Table S1 –** DSM diagnostic labels of patients with bipolar disorder.

| DSM-5 | n |
|---|---|
| 296.40 | 8 |
| 296.41 | 1 |
| 296.42 | 5 |
| 296.45 | 3 |
| 296.46 | 4 |
| 296.51 | 6 |
| 296.52 | 7 |
| 296.55 | 1 |
| 296.62 | 2 |
| 296.89 | 7 |
| 296.02 | 1 |
| 296.43 | 1 |

**Table S2 -** Demographics of schizophrenia patients (SZ), patients with bipolar disorder (BP) and controls (ctrl) who performed the cognitive tasks.

| | Degraded continuous performance test | | | Wisconsin card sorting test | | | Verbal fluency test | | |
|---|---|---|---|---|---|---|---|---|---|
| | SZ | ctrl | BP | SZ | ctrl | BP | SZ | ctrl | BP |
| N | 120 | 93 | 45 | 122 | 94 | 45 | 52 | 66 | 24 |
| Gender (F/M) | 17/103 | 46/47 | 31/14 | 17/105 | 47/47 | 31/14 | 6/46 | 4/62 | 0/21 |
| Age | 36.3 ± 0.8 | 35.1 ± 0.9 | 35.2 ± 1.5 | 36.2 ± 0.8 | 35.2 ± 0.9 | 35.8 ± 1.5 | 38.2 ± 1.1 | 35.2 ± 0.9 | 36.0 ± 1.8 |
| Education (years) | 13.4 ± 0.2 | 15.2 ± 0.3 | 14.3 ± 0.3 | 13.3 ± 0.2 | 15.2 ± 0.3 | 14.3 ± 0.3 | 13.5 ± 0.3 | 15.2 ± 0.3 | 14.6 ± 0.5 |
| Handedness (L/R)[a] | 6/114 | 6/87 | 0/42 | 6/116 | 6/88 | 0/42 | 6/46 | 4/63 | 0/20 |
| Visual acuity | 1.4 ± 0.03 | 1.6 ± 0.04 | 1.3 ± 0.1 | 1.4 ± 0.03 | 1.6 ± 0.04 | 1.3 ± 0.1 | 1.4 ± 0.05 | 1.6 ± 0.05 | 1.2 ± 0.1 |
| Illness duration (years) | 12.0 ± 0.7 | | 11.6 ± 1.3 | 12.0 ± 0.7 | | 11.7 ± 1.3 | 13.5 ± 1.1 | | 11.3 ± 1.5 |
| SANS | 10.6 ± 0.5 | | | 10.5 ± 0.5 | | | 9.9 ± 0.7 | | |
| SAPS | 9.8 ± 0.7 | | | 9.7 ± 0.6 | | | 9.9 ± 1.4 | | |
| BPRS[b] | 33.0 ± 0.5 | | 31.4 ± 1.0 | 33.0 ± 0.5 | | 31.8 ± 1.0 | 33.0 ± 0.5 | | 33.4 ± 1.4 |
| CPZ equivalent[c] | 589.3 ± 36.7 | | 380.9 ± 53.1 | 586.0 ± 36.4 | | 367.4 ± 53.8 | 588.9 ± 57.2 | | 268.6 ± 34.3 |

Abbreviations: SANS, Scale for the Assessment of Negative Symptoms; SAPS, Scale for the Assessment of Positive Symptoms; BPRS, Brief Psychiatric Rating Scale; CPZ, Chlorpromazine equivalents.

[a] Data from 3 BP were missing for all the three cognitive tasks.

[b] Only 31/123 SZ were considered for all the three cognitive tasks.

[c] Patients receiving medication: CPT: 108/120 SZ, 35/45 BP; WCST: 109/122 SZ, 35/45 BP; VFT: 47/52 SZ, 19/24 BP.

**Table S3 –** GFP amplitudes at the peak latencies (± SD), expressed in μV, for each group and each condition. Results are graphically shown in Fig. 3B.

| Mean ± SD (peak latency) | SZ | ctrl | BP |
|---|---|---|---|
| Vernier Only | 1.8 ± 0.1 (209 ms) | 2.9 ± 0.1 (197 ms) | 2.1 ± 0.2 (213 ms) |
| Long SOA | 1.8 ± 0.1 (209 ms) | 2.9 ± 0.1 (197 ms) | 2.2 ± 0.3 (209 ms) |
| Short SOA | 2.1 ± 0.1 (195 ms) | 3.4 ± 0.2 (191 ms) | 2.1 ± 0.3 (189 ms) |
| Mask Only | 1.7 ± 0.1 (146 ms) | 2.0 ± 0.1 (170 ms) | 1.7 ± 0.2 (145 ms) |

**Table S4 -** Supplementary statistical analysis for the visual backward masking performance: adaptive masking experiment (5- and 25-elements masks) and EEG experiment.

| **5-elements and 25-elements masks (SOAs)** | | | |
|---|---|---|---|
| **mask** | $F$(1,235)=14.796, $\eta^2$=.011, $P$<.001 | | |
| **mask * group** | $F$(2,235)=1.117, $\eta^2$=.002, $P$=.329 | | |
| **mask * gender** | $F$(1,235)=.394, $\eta^2$<.001, $P$=.531 | | |
| **mask * visual acuity** | $F$(1,235)=.394, $\eta^2$<.001, $P$=.912 | | |
| **mask * education** | $F$(1,235)=3.894, $\eta^2$=.003, $P$=.050 | | |
| **mask * group * gender** | $F$(2,235)=.087, $\eta^2$<.001, $P$=.917 | | |
| **group** | $F$(2,235)=22.916, $\eta^2$=.159, $P$<.001 | | |
| post-hoc ANCOVA | **ctrl vs SZ** | **ctrl vs BP** | **SZ vs BP** |
| | $F$(1,404)=52.332, $\eta^2$=.112, $P_{holm}$<.001, d=-.710 | $F$(1,258)=30.570, $\eta^2$=.103, $P_{holm}$<.001, d=-.678 | $F$(1,292)=1.645, $\eta^2$=.006, $P_{holm}$=.201, d=.155 |
| **gender** | $F$(1,235)=3.205, $\eta^2$=.011, $P$=.075 | | |
| **visual acuity** | $F$(1,235)=2.348, $\eta^2$=.008, $P$=.127 | | |
| **education** | $F$(1,235)=1.349, $\eta^2$=.005, $P$=.247 | | |
| **group * gender** | $F$(2,235)=.253, $\eta^2$=.002, $P$=.777 | | |
| **Percent correct in the EEG experiment** | | Greenhouse-Geisser as assumption of sphericity is violated | |
| **condition** | $F$(1.810,403.525)=29.982, $\eta^2$=.049, $P$<.001 | | |
| **condition * group** | $F$(3.619,403.525)=16.881, $\eta^2$=.055, $P$<.001 | post-hoc tests were reported in the supplementary Table S6 | |
| **condition * gender** | $F$(1.810,403.525)=7.845, $\eta^2$=.013, $P$<.001 | | |
| **condition * education** | $F$(1.810,403.525)=.322, $\eta^2$<.001, $P$=.703 | | |
| **condition * visual acuity** | $F$(1.810,403.525)=2.874, $\eta^2$=.005, $P$=.063 | | |
| **condition * group * gender** | $F$(3.619,403.525)=2.249, $\eta^2$=.007, $P$=.070 | | |
| **group** | $F$(2,223)=21.768, $\eta^2$=.153, $P$<.001 | | |
| **gender** | $F$(1,223)=7.614, $\eta^2$=.027, $P$=.006 | | |
| **education** | $F$(1,223)=.360, $\eta^2$=.001, $P$=.549 | | |
| **visual acuity** | $F$(1,223)=4.393, $\eta^2$=.015, $P$=.037 | | |
| **group * gender** | $F$(2,223)=3.224, $\eta^2$=.023, $P$=.042 | | |

Since the three study groups differed in terms of gender, education, and VA, a supplementary statistical analysis for the backward masking task was conducted using gender as a factor and education and VA as covariates. rm-ANOVA to test SOAs was followed by three ANCOVAs to test for group effect (SOAs measured were combined in this case). Eta squares were transformed into Cohen's d with the aid of https://www.psychometrica.de/effect_size.html#transform, and direction was inferred from the data. Regarding the performance in the EEG experiment, rm-ANOVA was conducted. Post-hoc ANCOVA for the condition * group interaction were reported in supplementary Table S6.

***Comment on the results:*** In the adaptive task, a main effect of mask, group, and an interaction mask * education were found. Post-hoc ANCOVAs for the group effect showed that SZ and BP performed significantly worse than controls, as it was the case for the main analysis. The absolute effect sizes were larger in the main analysis for the significant comparisons (main: ctrl-SZ: d=.871; ctrl-BP: d=.794, SZ-BP: d=.092; supplementary: ctrl-SZ: d=.710; ctrl-BP: d=.678, SZ-BP: d=.155). The fact that education, visual acuity and gender did not influenced the results is in agreement with previous studies, where the inclusion of the covariates did not influence VBM performance (2–4). In the EEG task, besides the condition, condition x group, and group effects, the interactions condition x gender, and group x gender, as well as a main effect of gender and visual acuity, were found. Results of the post-hoc tests for the condition x group effect (Table S6) showed that findings are comparable to the main analysis (i.e., significant differences between SZ and controls for the three conditions with the target vernier). In addition, a significant difference between SZ and BP was found for the EEG performance in the Short SOA condition.

**Table S5 -** Supplementary statistical analysis for the GFP N1 peak measured in the EEG experiment.

| **N1 peak (~200ms)** | Greenhouse-Geisser as assumption of sphericity is violated | |
|---|---|---|
| **condition** | $F$(2.105,469.463)=1.409, $\eta^2$=.001, $P$=.245 | |
| **condition * group** | $F$(4.210,469.463)=6.122, $\eta^2$=.011, $P$<.001 | post-hoc tests were reported in the supplementary Table S6 |
| **condition * gender** | $F$(2.105,469.463)=2.084, $\eta^2$=.002, $P$=.123 | |
| **condition * education** | $F$(2.105,469.463)=.236, $\eta^2$<.001, $P$=.801 | |
| **condition * VA** | $F$(2.105,469.463)=.376, $\eta^2$<.001, $P$=.698 | |
| **condition * group * gender** | $F$(4.210,469.463)=.393, $\eta^2$<.001, $P$=.823 | |

| | |
|---|---|
| **group** | $F$(2,223)=10.453, $\eta^2$=.084, $P$<.001 |
| **gender** | $F$(1,223)=1.879, $\eta^2$=.008, $P$=.172 |
| **education** | $F$(1,223)=.883, $\eta^2$=.004, $P$=.348 |
| **visual acuity** | $F$(1,223)=.732, $\eta^2$=.003, $P$=.393 |
| **group * gender** | $F$(2,223)=.888, $\eta^2$=.007, $P$=.413 |

rm-ANOVA was performed for the peak amplitude of the GFP grand average (N1 component), for each group and each condition, using gender as a factor and visual acuity and education as covariates. Post-hoc ANCOVA for the condition * group interaction were reported in supplementary Table S6.

***Comment on the results***: Only condition x group and group main effect were significant. Thus, no additional effects were added nor by gender nor the covariates. Post-hoc ANCOVAs for the condition x group interaction (Table S6) showed the same significant results as in the main analysis, although the effect sizes for the significant comparisons were reduced (ctrl-SZ main vs. supplementary: VO, d=.877 vs. d=.557; LSOA, d=.935 vs. d=.585; SSOA, d=.901 vs. d=.585; ctrl-BP for SSOA condition: d=.909 in main analysis, while d=.563 in the supplementary one). The marginal significances between controls and BP for the VO and Long SOA conditions found in the main analysis were no longer present when gender and covariates were taken into account.

### **Table S6 –** Complementary statistical analysis for the visual backward masking performance in the EEG experiment and for the GFP N1 peak.

| **Percent correct in the EEG experiment** | | | | |
|---|---|---|---|---|
| **condition * group** | $F$(3.622,403.891)=16.876, $\eta^2$=.056, $P$<.001 | | | |
| post-hoc ANCOVA | **ctrl vs SZ** | **ctrl vs BP** | **SZ vs BP** | **ANCOVA** |
| Vernier Only | $F$(1,209)=21.466, $\eta^2$=.091, $P$<.001, d=.633 | $F$(1,104)=.302, $\eta^2$=.002, $P$=.584, d=.090 | $F$(1,131)=4.490, $\eta^2$=.032, $P$=.170, d=-.364 | $F$(2,223)=11.442, $\eta^2$=.090, $P$<.001 |
| Long SOA | $F$(1,209)=29.433, $\eta^2$=.120, $P$<.001, d=.739 | $F$(1,104)=4.627, $\eta^2$=.034, $P$=.170, d=.375 | $F$(1,131)=2.232, $\eta^2$=.016, $P$=.414, d=-.255 | $F$(2,223)=15.021, $\eta^2$=.113, $P$<.001 |
| Short SOA | $F$(1,209)=49.446, $\eta^2$=.185, $P$<.001, d=.953 | $F$(1,104)=1.327, $\eta^2$=.011, $P$=.504, d=.211 | $F$(1,131)=8.770, $\eta^2$=.056, $P$=.024, d=-.487 | $F$(2,223)=24.599, $\eta^2$=.167, $P$<.001 |
| Mask Only | | | | $F$(2,223)=.495, $\eta^2$=.002, $P$=.482 |
| **N1 peak (~200ms)** | | | | |
| **condition * group** | $F$(4.210,469.463)=6.122, $\eta^2$=.011, $P$<.001 | | | |
| post-hoc ANCOVA | **ctrl vs SZ** | **ctrl vs BP** | **SZ vs BP** | **ANCOVA** |
| Vernier Only | $F$(1,209)=17.130, $\eta^2$=.072, $P$<.001, d=.557 | $F$(1,104)=3.666, $\eta^2$=.033, $P$=.290, d=.370 | $F$(1,131)=.106, $\eta^2$<.001, $P$=1.000, d=-.056 | $F$(2,223)=9.518, $\eta^2$=.077, $P$<.001 |
| Long SOA | $F$(1,209)=19.042, $\eta^2$=.079, $P$<.001, d=.586 | $F$(1,104)=3.282, $\eta^2$=.030, $P$=.292, d=.352 | $F$(1,131)=.339, $\eta^2$=.002, $P$=1.000, d=-.090 | $F$(2,223)=10.195, $\eta^2$=.082, $P$<.001 |
| Short SOA | $F$(1,209)=19.039, $\eta^2$=.079, $P$<.001, d=.586 | $F$(1,104)=7.584, $\eta^2$=.067, $P$=.042, d=.536 | $F$(1,131)=.601, $\eta^2$=.004, $P$=1.000, d=.127 | $F$(2,223)=12.592, $\eta^2$=.099, $P$<.001 |
| Mask Only | | | | $F$(2,223)=1.835, $\eta^2$=.016, $P$=.162 |

Eta square were transformed into Cohen's d with the aid of https://www.psychometrica.de/effect_size.html#transform, and direction was inferred from the data. P-values Bonferroni-Holm corrected for multiple comparisons.

### **Table S7 -** Pearson Correlation of BPRS and CPZ data on the VBM task results (adaptive and EEG).

| | **BPRS** | | **CPZ** | |
|---|---|---|---|---|
| | SZ | BP | SZ | BP |
| **Adaptive Experiment *** | | | | |
| SOA 5 | r(28)=.002, p=.993 | r(36)=-.098, p=.558 | r(109)=.080, p=.406 | r(36)=.205, p=.217 |
| SOA 25 | r(28)=.129, p=.497 | r(36)=-.011, p=.949 | r(109)=.189, p=.048 | r(36)=.190, p=.252 |

| | | | | |
|---|---|---|---|---|
| **EEG Experiment Behavior **** | | | | |
| Vernier Only | r(28)=-.370, p=.044 | r(14)=-.215, p=.423 | r(119)=-.286, p=.001 | r(14)=-.158, p=.560 |
| Long SOA | r(28)=-.412, p=.024 | r(14)=-.338, p=.201 | r(119)=-.235, p=.010 | r(14)=-.080, p=.770 |
| Short SOA | r(28)=-.115, p=.546 | r(16)=-.273, p=.307 | r(119)=-.276, p=.002 | r(14)=-.264, p=.323 |
| **EEG Experiment GFP Amplitude **** | | | | |
| Vernier Only | r(28)=-.123, p=.517 | r(14)=.284, p=.286 | r(119)=-.200, p=.028 | r(14)=-.151, p=.577 |
| Long SOA | r(28)=-.213, p=.258 | r(14)=.236, p=.380 | r(119)=-.230, p=.012 | r(14)=-.194, p=.471 |
| Short SOA | r(28)=-.078, p=.683 | r(14)=.220, p=.413 | r(119)=-.165, p=.070 | r(14)=-.214, p=.426 |
| Mask Only | r(28)=.090, p=.638 | r(16)=-.480, p=.061 | r(119)=.063, p=.496 | r(14)=-.164, p=.544 |

Abbreviations: SOA, Stimulus Onset Asynchrony; BPRS, Brief Psychiatric Rating Scale; CPZ, Chlorpromazine equivalents.
P-values are not corrected for multiple comparisons.
* BPRS: data from 31 SZ, 43 BP are included. CPZ: data from 122 SZ, 43 BP are included.
**BPRS: data from 30 SZ, 16 BP are included. CPZ: data from 121 SZ, 16 BP are included.

***Comment on the results***: For the SZ and BP groups, symptoms and CPZ equivalent were independently correlated with VBM performance and GFP N1 peak. SZ and BP had no significantly different overall symptoms (adaptive: d=-.268, EEG: d=-.059). SZ received significantly more medications than BP (adaptive: d=.517, EEG: d=1.173). Symptoms analysis gave only a weak correlation for the EEG task accuracy in SZ, for the Vernier Only and Long SOA conditions. Here, SZ with a lower BPRS values had higher accuracy. The fact that, generally, symptoms do not correlate with VBM outcomes may indicates that the task can discriminate all patients, independently of the severity of the disorder. Correlation of CPZ equivalent showed that SZ receiving less medication performed better in the SOA25 condition of the adaptive experiment, as well as in the three conditions with the target vernier of the EEG task. Additionally, SZ receiving less medication had a higher GFP N1 peak in the Vernier Only and Long SOA conditions. No significant correlation between CPZ and BP measures was found, probably due to a lack of statistical power.

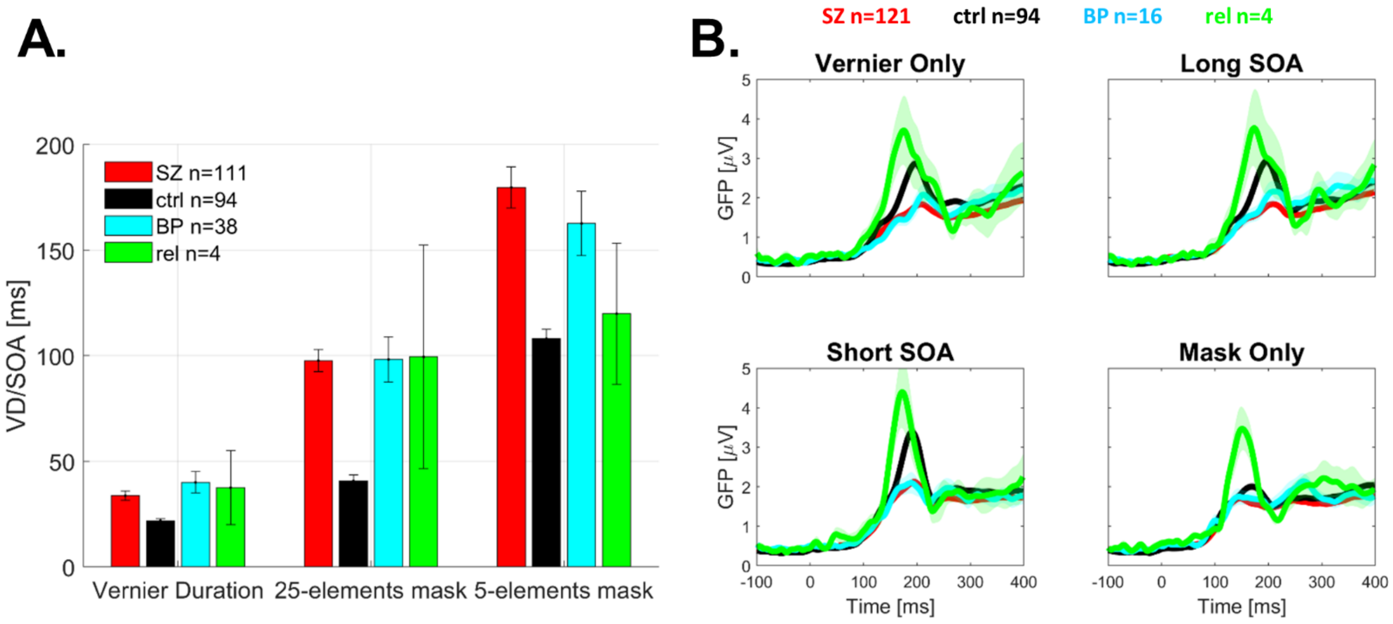


**Figure S1: Very preliminary results for the siblings of patients with bipolar disorder (green).** (A) Behavioral results of the adaptive experiment: VDs and stimulus onset asynchrony (SOA) for the two types of masks. (Note: SOA=VD+ISI, longer SOAs=stronger deficits). Error bars represent the standard error of the mean. (B) Group average global field power (GFP) time series in each condition.

Shaded areas indicate SEM. Sibling of BP (green) showed higher GFP amplitudes compared to controls (black), may suggesting a compensation mechanism, potentially similar to reported in da Cruz and colleagues (5).
